# Linking Activity in Human Superior Temporal Cortex to Perception of Noisy Audiovisual Speech

**DOI:** 10.1101/2020.04.02.021774

**Authors:** Johannes Rennig, Michael S Beauchamp

## Abstract

Regions of the human posterior superior temporal gyrus and sulcus (pSTG/S) respond to the visual mouth movements that constitute visual speech and the auditory vocalizations that constitute auditory speech. We hypothesized that these multisensory responses in pSTG/S underlie the observation that comprehension of noisy auditory speech is improved when it is accompanied by visual speech. To test this idea, we presented audiovisual sentences that contained either a clear auditory component or a noisy auditory component while measuring brain activity using BOLD fMRI. Participants reported the intelligibility of the speech on each trial with a button press. Perceptually, adding visual speech to noisy auditory sentences rendered them much more intelligible. Post-hoc trial sorting was used to examine brain activations during noisy sentences that were more or less intelligible, focusing on multisensory speech regions in the pSTG/S identified with an independent visual speech localizer. Univariate analysis showed that less intelligible noisy audiovisual sentences evoked a weaker BOLD response, while more intelligible sentences evoked a stronger BOLD response that was indistinguishable from clear sentences. To better understand these differences, we conducted a multivariate representational similarity analysis. The pattern of response for intelligible noisy audiovisual sentences was more similar to the pattern for clear sentences, while the response pattern for unintelligible noisy sentences was less similar. These results show that for both univariate and multivariate analyses, successful integration of visual and noisy auditory speech normalizes responses in pSTG/S, providing evidence that multisensory subregions of pSTG/S are responsible for the perceptual benefit of visual speech.

**Significance Statement:** Enabling social interactions, including the production and perception of speech, is a key function of the human brain. Speech perception is a complex computational problem that the brain solves using both visual information from the talker’s facial movements and auditory information from the talker’s voice. Visual speech information is particularly important under noisy listening conditions when auditory speech is difficult or impossible to understand alone Regions of the human cortex in posterior superior temporal lobe respond to the visual mouth movements that constitute visual speech and the auditory vocalizations that constitute auditory speech. We show that the pattern of activity in cortex reflects the successful multisensory integration of auditory and visual speech information in the service of perception.

## Introduction

Enabling social interactions, including the production and perception of speech, is a key function of the human brain. Speech perception is a complex computational problem that the brain solves using both visual information from the talker’s facial movements and auditory information from the talker’s voice. Visual speech information is particularly important under noisy listening conditions when auditory speech is difficult or impossible to understand alone (reviewed in 1).

The neural substrates for the integration of auditory and visual speech are a subject of active investigation (2). In non-human primates, recordings from single neurons in pSTG/S respond to both auditory and visual social communication signals (3-5). In humans, small populations of neurons in pSTG/S recorded with intracranial electrodes respond to both auditory and visual speech (6, 7). While the idea that pSTG/S integrates visual speech information with noisy auditory speech in the service of comprehension seems reasonable, it is supported by only limited empirical evidence. In a recent study, repetitive transcranial magnetic stimulation (rTMS) was used to disrupt processing of pSTG/S during perception of noisy audiovisual sentences (8). rTMS resulted in small but significant decreases (between one and two dB) in the ability to understand noisy speech. However, a BOLD fMRI study failed to find evidence that pSTG/S integrates noisy auditory and visual speech. Bishop and Miller presented noisy audiovisual syllables and used *post hoc* sorting to separate trials into those that were understood and those that were not (9). Differential BOLD fMRI responses to the two types of trials was observed in seventeen different brain areas, but pSTG/S was not of them.

As in most published neuroimaging studies of noisy speech perception, Bishop and Miller performed a univariate analysis, examining the response amplitude in *individual* voxels and regions of interest. However, in many circumstances, multivariate analyses of the pattern of activity across *multiple* voxels reveals information hidden from univariate analyses (10). Another widely-used technique applied by Bishop and Miller was the use of a volumetric group analysis in which each participant was aligned to a template brain and analysis was conducted at the group level. While pSTG/S was classified as a single cytoarchitectonic area by Brodmann (BA 21), modern studies suggest that it contains a dozen or more functional subdivisions with boundaries that vary between individuals (11). Volumetric group analysis ignores this anatomical variability and assumes that a given coordinate in standard space is functionally equivalent across participants, sometimes leading to incorrect inferences (12).

We have recently developed a localizer that identifies regions important for processing audiovisual speech in the pSTG/S of individual participants (13). Inspired by earlier work (14, 15), the localizer contrasts responses to silent videos of actors making mouth movements with silent videos of actors making eye movements. Even though the localizer contains only unisensory visual stimuli, it successfully locates regions of pSTG/S that respond to voices and prefer voices to environmental sounds (16). A follow-up study demonstrated that mouth-preferring and eye-preferring subregions of pSTG/S could be identified during free viewing of talking faces by measuring eye movements made by participants in the MR scanner (17).

We applied this localizer to re-examine the relationship between comprehension of noisy audiovisual speech and BOLD responses in pSTG/S. Because the localizer identifies a population of voxels likely to be important for multisensory speech processing, it allows for the application of multivariate analysis techniques that compare the response pattern across conditions. Unlike volumetric group analyses, the localizer compares “apples to apples” since it identifies functionally equivalent regions across participants.

## Methods

Twenty-two healthy right-handed participants (14 females, mean age 25, range 18 - 34) with normal or corrected to normal vision and normal hearing provided written informed consent under an experimental protocol approved by the Committee for the Protection of Human Subjects of the Baylor College of Medicine, Houston, TX.

### Localizer fMRI Experiment

The localizer fMRI experiment was adapted from (Zhu and Beauchamp, 2017). The visual stimuli consisted of silent 2-second movies of two different actors making a variety of mouth movements and eye movements. Following each video, there was a 1-second response period during which participants pressed a button to identify the actor in the video. Stimuli were presented in a block design, with each block containing movies of a single type, either mouth movements or eye movements. Each 30-second block contained ten 3-second trials (2-second stimulus + 1-second response) followed by 10 seconds of fixation baseline. Three mouth and three eye blocks were presented alternately during each scan series, for a total duration of 240 seconds; two localizer scan series were collected for each participant.

### Main fMRI Experiment

The main fMRI experiment used an event-related design. Each 6-second trial consisted of the presentation of a 3-second recording of a sentence, followed by a 3-second response period. The sentences were recorded from a single male talker and were presented in five different formats: audiovisual (AV, video and clear audio), auditory (A, only clear audio), visual (V, only video), noisy audiovisual (AnV, video and noisy audio) and noisy auditory versions (An, only noisy audio) (Figure 1). To create noisy sentences, the audio recordings were combined with pink noise at a signal-to-noise ratio (SNR) of -16 dB. Pink noise, defined as noise with decreasing energy at increasing frequency, is less aversive than white noise and is commonly used in studies of auditory function. The sentence stimuli have been used in previous behavioral studies (Van Engen et al. 2017; Rennig et al. 2018).

**Figure 1:**
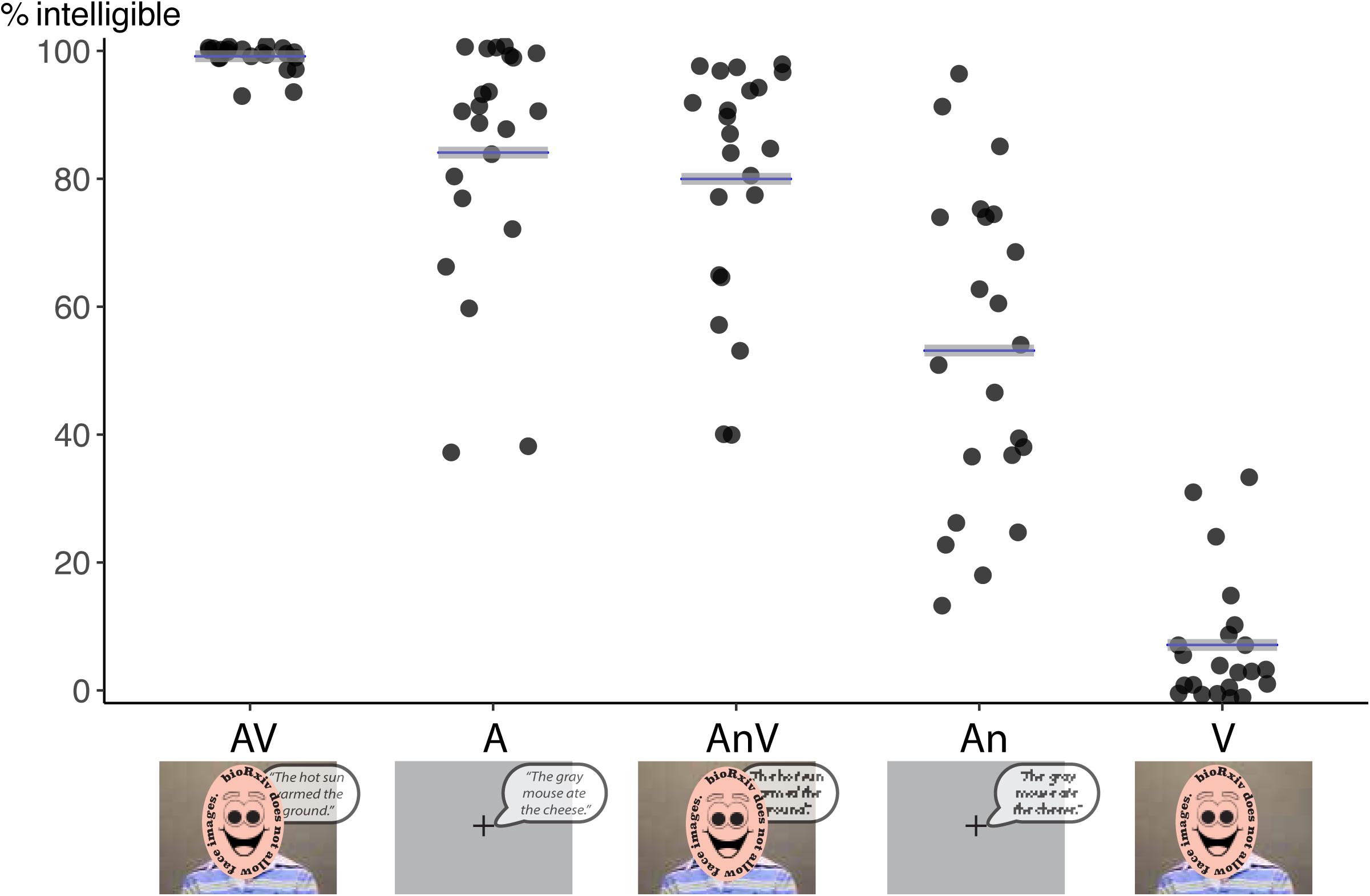
Behavioral responses to the five different types of sentences. After presentation of each sentence in the MR scanner, participants reported with a button press whether or not the sentence was intelligible. Black dots represent the percent of sentences rated as intelligible by a single participant; gray lines represent the mean across participants. The stimuli consisted of sentences presented in a clear audiovisual format (AV), auditory-only format (A), noisy-auditory with visual (AnV), noisy-auditory-only (An), or visual-only (V). Images show the visual stimulus, consisting of a video of a talking face (AV, AnV, V) or a blank screen with fixation crosshairs (A, An). Speech bubbles represent the auditory stimulus, degraded font illustrates auditory speech with added pink noise (AnV, An).

Four scan series were collected for each participant. Each series had total duration of 300 seconds and contained 40 sentence trials (8 from each of the 5 different formats) and 60 seconds of fixation baseline in a pseudo-random optimal sequence generated by the program *optseq2* (Dale, 1999; https://surfer.nmr.mgh.harvard.edu/optseq).

### MRI Acquisition

Participants were scanned in a 3 tesla Siemens Trio MRI scanner equipped with a 32-channel head coil at Baylor College of Medicine’s Core for Advanced MRI. Stimuli were presented using Matlab (The Mathworks, Inc., Natick, MA, USA) with the Psychophysics Toolbox extensions (Brainard 1997; Pelli 1997). Visual stimuli were presented on an MR compatible screen (BOLDscreen32, Cambridge Research Systems, Rochester, UK) placed behind the bore of the scanner and viewed through a mirror. Auditory stimuli were presented using high-fidelity MR compatible headphones (Sensimetrics, Malden, MA, USA). Behavioral responses were collected using a fiber-optic button response pad (Current Designs, Haverford, PA, USA) and eye movements were recorded during scanning using the Eye Link 1000 (SR Research Ltd., Ottawa, Ontario, Canada) with a sampling rate of 500 Hz.

Two T1-weighted MP-RAGE anatomical volumes and six EPI scan series (two localizer and four for the main experiment) were collected from each participant. EPI data was acquired using a multi-slice echo planar imaging sequence (Setsompop et al. 2012): TR = 1500 ms, TE = 30 ms, flip angle = 72°, in-plane resolution of 2 × 2 mm, 69 2 mm axial slices, multiband factor: 3, GRAPPA factor: 2.

### Perceptual Ratings

After each sentence presentation in the main fMRI experiment, participants rated intelligibility using a three-alternative forced choice (to minimize perceptual learning, sentences were not repeated within participants). The rating choices were: “understood everything” (all words in the sentence understood); “understood something” (at least one word in the sentence); “understood nothing” (no words in the sentence). To ensure sufficient trial numbers for *post-hoc* trial sorting, the “everything” and “something” choices were grouped together, resulting in two types of perceptually-sorted trials: “yes—some or all of the sentence intelligible” (Y) and “not at all intelligible” (N). This grouping was necessary because there were few “understood everything” responses (8% in the auditory-only noisy condition).

### fMRI analysis

fMRI analysis was conducted using AFNI (Cox 1996). Preprocessing consisted of slice-timing correction and motion correction by maximizing local Pearson correlation (Saad et al. 2009). The generalized linear model (GLM) regressors of no interest included a third order polynomial (to model baseline fluctuations) and six motion parameters (roll, pitch, yaw; linear movement into x-, y-, z-directions). For the localizer fMRI experiment, there were two regressors of interest, one for all blocks containing mouth movements and one for all blocks containing eye movements. For the main fMRI experiment, the initial GLM contained five regressors of interest, one for each sentence type (A, An, AV, AnV, V). In a second analysis, the noisy sentences were *post-hoc* sorted by perceptual ratings into sentences that were intelligible (Y) or were not intelligible (N), producing a GLM with seven regressors of interest (A, An-Y, An-N, AV, AnV-Y, AnV-N, V).

The regressors of interest were created by convolving the time of each stimulus presentation with an incomplete gamma function using the BLOCK() function in the AFNI program *3dDeconvolve*. In order to estimate the time course of the BOLD response, an separate GLM was constructed using the TENT() function, with a separate regressor for each time point of the response in the window from zero to fifteen seconds after stimulus onset.

### ROI construction

For every participant, we created a cortical surface model from two repetitions of a T1-weighted image with FreeSurfer (Dale et al. 1999; Fischl et al. 2002) and manipulated it with SUMA (Argall et al. 2006). As the anterior and posterior temporal cortex are functionally distinct, posterior temporal cortex was defined as cortex in the posterior half of the Freesurfer-defined superior temporal gyrus, superior temporal sulcus, and middle temporal gyrus labels (Destrieux et al. 2010). Across subjects, the average location of the ROI midpoint was y = -25 ± 1.5 mm (left hemisphere) and y = -24 ± 0.8 mm (right hemisphere); co-ordinates in MNI standard space (N27). pSTG/S mouth ROIs were created by finding voxels in the posterior temporal anatomical ROI that showed a significant overall omnibus *F*-test (*F* > 5, *q* < 0.0001, false discovery rate corrected) and a significant preference for mouth compared with eye videos in the localizer fMRI experiment (*q* < 0.05). pSTG/S eye ROIs were defined as posterior temporal voxels that showed the reverse preference (eye videos > mouth videos, *q* < 0.05). Sample ROIs from 4 participants are shown in Figure S1.

### Mixed-effects models

Analysis across participants was conducted using linear mixed-effects (LME) models created with the *lme4* package in R (Bates et al. 2015). For each participant, the average BOLD fMRI response in each ROI in each hemisphere was calculated. To determine if hemisphere was an important predictor of the BOLD response, we fit two models. The first model used three fixed factors: ROI (mouth-preferring cortex, eye-preferring cortex); stimulus condition (A, An, AV, AnV, V) and hemisphere (left, right). The second model used only two fixed factors: ROI and stimulus condition. Participant was included as a random factor in both models. The two models were compared using the Bayesian information criterion (BIC), with a BIC difference between models of 10 or greater considered decisive (18). Since the model without the hemisphere factor explained the data better (BIC difference = 46) all analyses reported in the manuscript use the second model, without the hemisphere factor.

For each statistical test, the degrees of freedom, *t* value, and *p* value were calculated according to the Satterthwaite approximation using the *lmerTest* package (Kuznetsova et al. 2015) and ANOVA-like tests (Type II Wald chi square test resulting in *χ*^*2*^ and *p* values) were calculated using the *Anova* function of the *car* package.

### Multivariate analysis

Multivariate analysis was performed within individual participant ROIs. For a given condition, the multivariate response pattern was defined as the percent signal change in each voxel within the ROI evoked by that condition. Pair-wise comparisons between conditions were calculated as the simple linear correlation between the patterns evoked by each condition (19). It should be noted that unlike some analyses (e.g. 20) we did not subtract the mean value across conditions within each voxel before performing the correlation. This means that all reported pattern correlations are positive, reflecting the positive BOLD response to sentences in the pSTG/S.

## Results

### Perceptual Data

In the main experiment, participants were presented with 5 different types of sentences in the MR scanner (Fig. 1). Participants rated each sentence as intelligible (some or every word in the sentence understood) or unintelligible (no words understood). Consistent difference in intelligibility across conditions were observed (Figure 1). Audiovisual sentences (AV) were the most intelligible (99% of sentences rated intelligible), followed by clear auditory-only sentences (A, 84%), audiovisual sentences with pink noise added to the auditory track (AnV, 80%), auditory-only noisy sentences (An, 53%), and visual-only sentences (V, 10%). It should be noted that the echo-planar pulse sequence used for MR acquisition created a moderate degree of auditory noise that was equivalent across conditions.

A linear mixed-effects model with factors Speech Type (A, AV) and Presentation Type (clear, noisy) showed significant main effect for Speech Type (*χ*^*2*^_(1)_ = 33, *p* = 9.1 × 10^−9^) and Presentation Type (*χ*^*2*^_(1)_ = 47, *p* = 3.1 × 10^−12^) without a significant interaction (*χ*^*2*^_(1)_ = 2.6, *p* = 0.12). The most informative pair-wise comparisons were between conditions with and without visual speech at the same level of auditory noise. For speech with added pink noise, visual speech improved intelligibility by 30% (AnV *vs*. An, 83% *vs*. 53%, *χ*^*2*^_(1)_ = 67, *p* = 2.7 × 10^−16^). For clear speech, the improvement was 15% (99% vs. 84%, AV *vs*. A, *χ*^*2*^_(1)_ = 15, *p* = 0.0001).

### Functional Localizer: Identification of mouth-preferring cortex in pSTG/S

In each participant, the localizer fMRI experiment identified regions of the posterior temporal cortex that preferred visually-presented silent mouth movements to silent eye movements. Figure 2A shows an illustration of the mouth-preferring ROI on a group-averaged cortical surface (see Fig. S1 for results in 8 example hemispheres).

**Figure 2:**
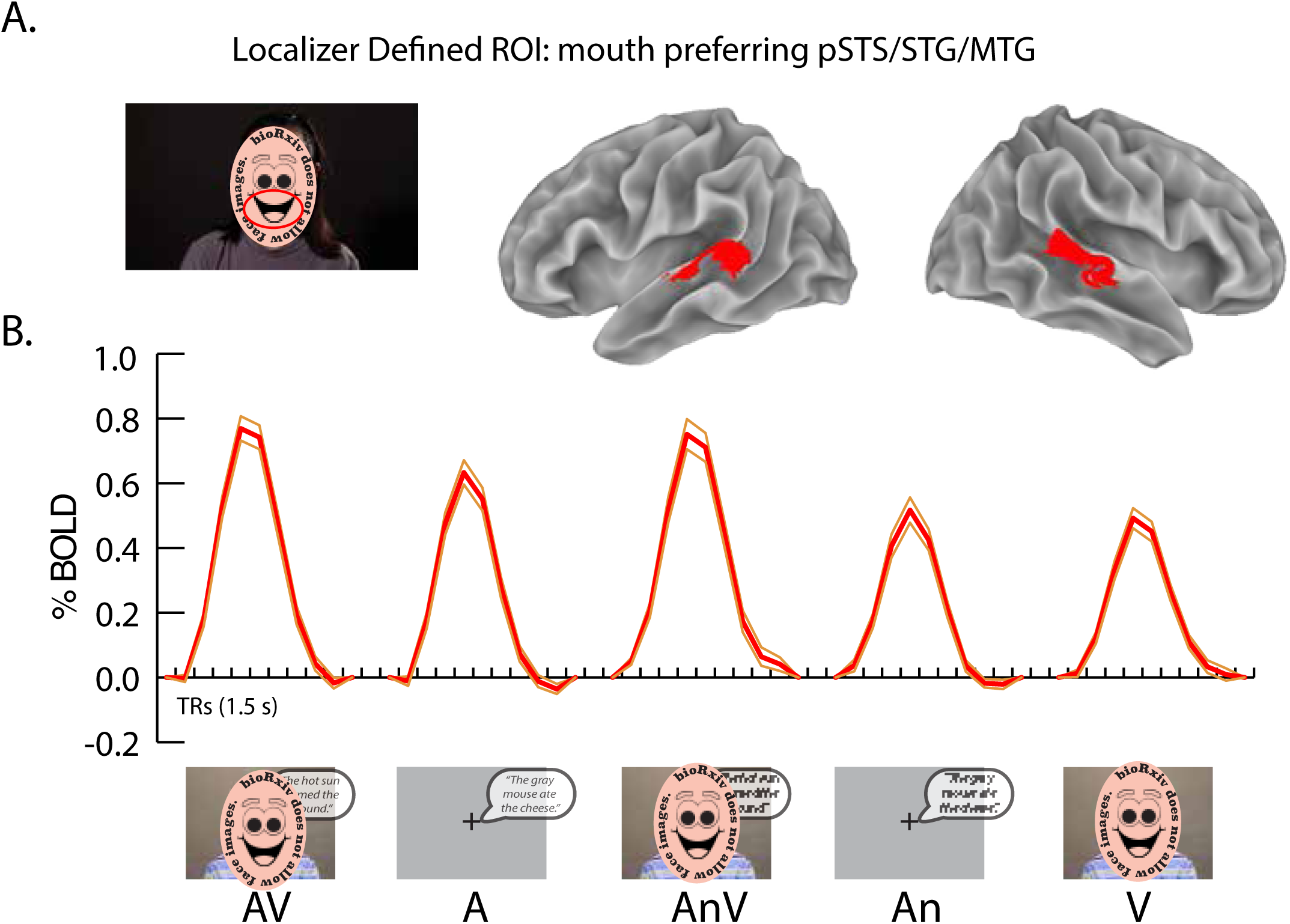
**A**. In the localizer fMRI experiment, silent videos of facial eye and mouth movements were presented. Image shows still frame from silent video of mouth movements, red ellipse highlights mouth movement. In each individual participant, mouth-preferring regions in the left and right pSTG/S were identified. For visualization only, a group map of mouth-preferring pSTG/S cortex was created (red color; Fig. S1 shows ROIs in individual participants). **B**. For each participant, the time course of the BOLD fMRI responses to the five different sentence types was calculated within the mouth-preferring ROI. Thick red lines show mean across participants, thin orange lines show standard error of the mean.

### BOLD Response to Clear and Noisy Sentences

We examined the response of mouth-preferring pSTG/S to the 5 types of sentences presented in the main experiment (Fig. 2B). AV sentences evoked the strongest response, significantly greater than either of the component unisensory conditions (AV *vs*. V: 0.75% vs. 0.46%; *χ*^*2*^_(1)_ = 64, *p* = 1.2 × 10^−15^; AV *vs*. A: 0.75% *vs*. 0.60%; *χ*^*2*^_(1)_ = 21, *p* = 4.4 × 10^−6^). For the unisensory sentences, A sentences evoked a larger response than V sentences (A *vs*. V: 0.60% vs. 0.46%; *χ*^*2*^_(1)_ = 14, *p* = 1.6 × 10^−4^).

Adding auditory noise significantly reduced the response for auditory-only sentences (A *vs*. An: 0.60% *vs*. 0.48%; *χ*^*2*^_(1)_ = 18, *p* = 2.8 × 10^−5^) but not audiovisual sentences (AV *vs*. AnV: 0.75% *vs*. 0.74%; *χ*^*2*^_(1)_ = 0.16, *p* = 0.69).

### BOLD Response to Noisy Sentences, Sorted by Perception

If the pSTG/S is the neural locus for the perceptual benefit of visual speech, we might expect to find a relationship between BOLD responses in the pSTG/S and sentence intelligibility. More effective integration of visual and auditory speech might lead to greater neural responses (higher BOLD signal) and improved perception. To examine this relationship, we applied the procedure of *post-hoc* sorting of fMRI data by perception. For each stimulus category, sentences rated as intelligible were placed in one bin and sentences rated as unintelligible were placed in the other (Figure 3). To minimize any confounding effects of stimulus, the initial analysis was only conducted within stimulus category, so that stimulus factors (such as the presence or absence of added noise) were equivalent between bins.

**Figure 3:**
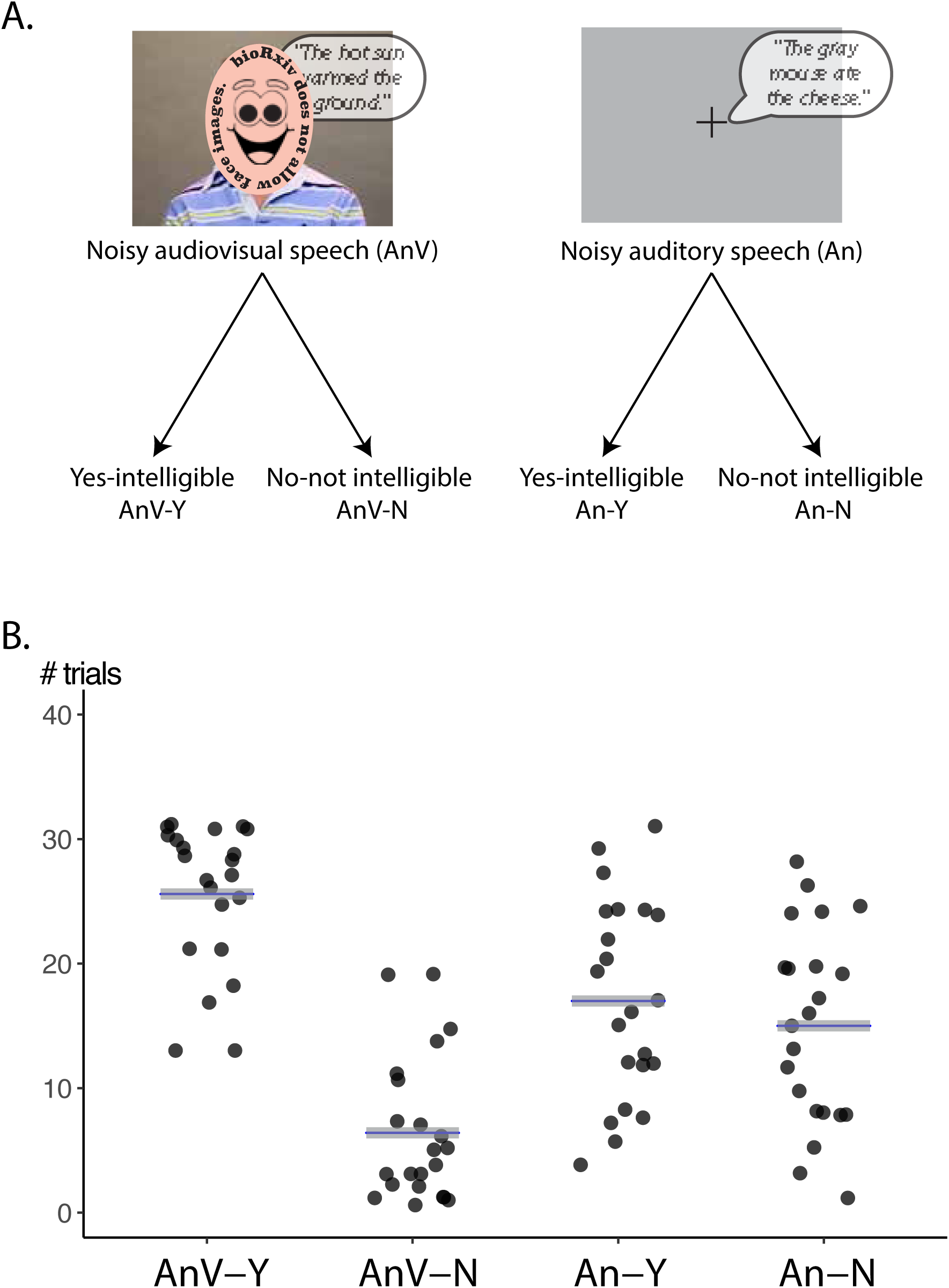
**A**. Noisy sentences (AnV and An) were sorted into two categories based on behavioral responses, either intelligible (Y) or not intelligible (N), resulting in four trial types. **B**. The number of trials for each trial type. Black dots represent individual participants; gray lines represent the mean across participants.

The response to noisy auditory sentences rated as intelligible was significantly greater than the response to auditory-noisy sentences rated as unintelligible, An-Y *vs*. An-N, 0.50% *vs*. 0.44%; χ^*2*^_(1)_ = 4.3, *p* = 0.038 (Figure 4A). Similarly, the response to noisy audiovisual sentences rated as intelligible was significantly greater than the response to noisy audiovisual sentences rated as unintelligible, AnV-Y *vs*. AnV-N 0.74% *vs*. 0.62%, *χ*^*2*^_(1)_ = 5.53, *p* = 0.019 (Figure 4B). Overall, responses to clear auditory sentences were greater than responses to either intelligible or unintelligible noisy sentences (0.60 vs. 0.50%; *χ*^*2*^_(1)_ = 8.12, *p* = 0.004; 0.60% *vs*. 0.44%; *χ*^*2*^_(1)_ = 22.98, *p* = 1.6 × 10^−6^) while responses to clear audiovisual sentences were greater than responses to unintelligible noisy sentences (0.75% *vs*. 0.62%; *χ*^*2*^_(1)_ = 6.41, *p* = 0.011) but similar to those for intelligible noisy audiovisual sentences (0.75% *vs*. 0.74%; *χ*^*2*^_(1)_ = 0.00, *p* = 0.959).

**Figure 4:**
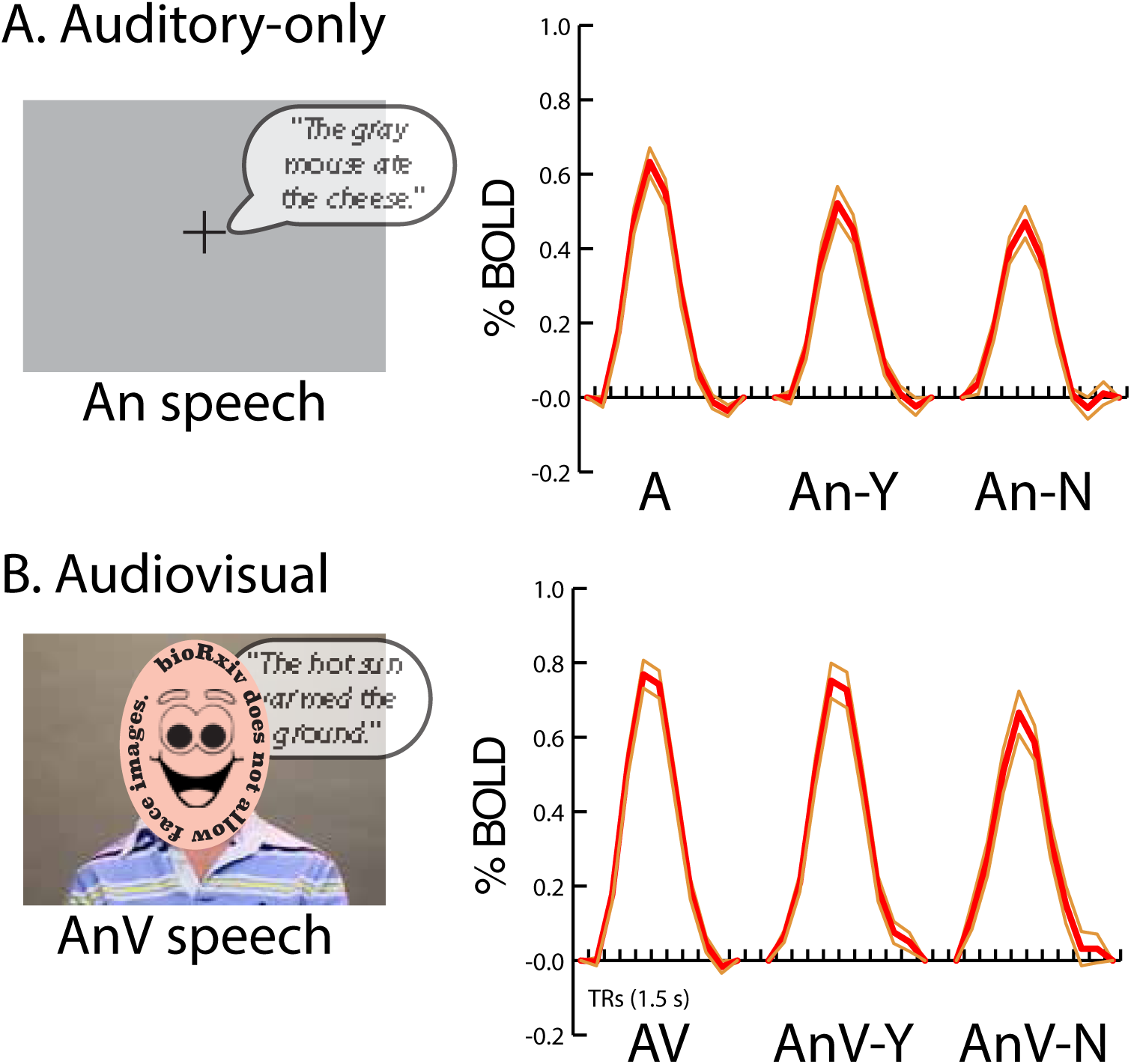
**A**. The time course of the BOLD fMRI response to noisy auditory speech rated as intelligible (An-Y) and noisy auditory speech rated as not intelligible (An-N) in mouth-preferring pSTG/S was calculated (response to A speech shown for reference). Thick red lines show mean across participants, thin orange lines show standard error of the mean. **B**. The time course of the BOLD fMRI response to noisy audiovisual speech rated as intelligible (AnV-Y) and noisy audiovisual speech rated as not intelligible (AnV-N). Response to AV speech shown for reference.

### Multivariate analysis

Next, we analyzed the multivariate response pattern in pSTG/S. For each pair of conditions, we correlated the response pattern of voxels within each participant’s mouth-preferring pSTG/S ROI, and then averaged across participants to calculate a mean *r*. Figure 5A shows the complete matrix of *r*-values between all pair-wise conditions.

**Figure 5:**
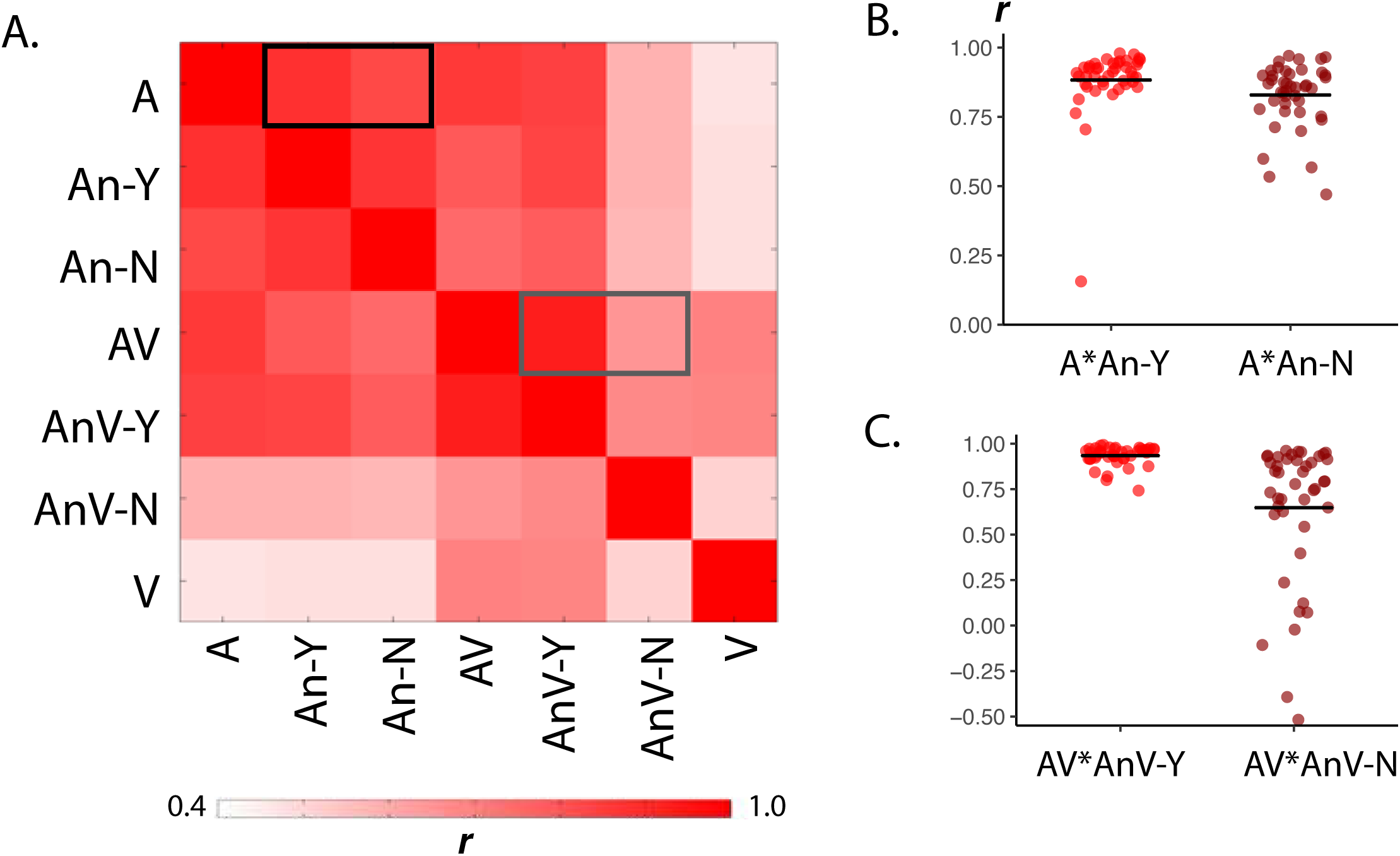
**A**. Average correlation matrix between all speech conditions in the mouth-preferring pSTG/S. For each participant, the activation patterns were correlated between every pair of conditions to create a matrix of correlation values. These matrices were then averaged across all participants. The colors indicate correlation strength (Pearson’s *r* coefficient). The black box highlights individual participant pairwise correlations shown in panel B. The gray box highlights individual participant pairwise correlations shown in panel C. **B**. Individual participant pattern correlations within mouth-preferring pSTG/S. Each dot represents the r coefficient of an individual participant, the black line represents the group mean (equal to the value shown in panel A). Left points (red) show pattern correlation between clear auditory (A) and noisy auditory speech rated as intelligible (An-Y). Right points (brown) show pattern correlation between clear auditory (A) and noisy auditory speech rated as not intelligible (An-N). **C**. Left points show pattern correlation between clear audiovisual (AV) and noisy audiovisual speech rated as intelligible (AnV-Y). Right points show pattern correlation between clear audiovisual (AV) and noisy audiovisual speech rated as not intelligible (AnV-N).

The lowest correlations were between visual-only and auditory-only sentences, the two conditions that differed the most in both sensory content and intelligibility. The average correlation between the response patterns for visual-only speech (which always received the lowest intelligibility ratings) and auditory-only speech (which always received high intelligibility ratings) was *r*_A*V_ = 0.47.

In contrast, there was much higher correlation between sentences with similar sensory content. Critically, intelligibility also influenced pattern correlation values, even when sensory factors were kept constant. There was a high correlation between the pSTG/S response pattern to clear auditory speech and noisy auditory sentences rated as intelligible, *r*_A*An-Y_ = 0.88. The correlation was significantly weaker for unintelligible noisy auditory sentences, *r*_A*An-N_ = 0.83; 0.88 *vs*. 0.83, *χ*^*2*^_(1)_ = 7.13, *p* = 0.0076 (Figure 5B).

The influence of intelligibility on the response pattern was even more pronounced for audiovisual speech. There was very high similarity between the response patterns to clear audiovisual speech and noisy audiovisual sentences rated as intelligible, *r*_AV*AnV-Y_ = 0.93 and much less similarity between clear audiovisual and unintelligible noisy audiovisual, *r*_AV*AnV-N_ = 0.65, resulting in a highly significant difference, 0.93 *vs*. 0.65, *χ*^*2*^_(1)_ = 32.57, *p* = 1.1 × 10^−8^ (Figure 5C).

To compare the effect of intelligibility across conditions, all correlations were entered into a linear mixed effects model with two fixed effects, stimulus type (A and AV) and percept (intelligible-Y and intelligible-N) and a random effect of participant. There were significant main effects for stimulus type (*χ*^*2*^_(1)_ = 4.7, *p* = 0.030) and percept (*χ*^*2*^_(1)_ = 32, *p* = 1.2 × 10^−8^) and a significant interaction (*χ*^*2*^_(1)_ = 15, *p* = 1.0 × 10^−4^). The interaction was driven by the larger increase in pattern similarity with intelligibility for audiovisual sentences than auditory sentences.

### Control Analysis: Identification of eye-preferring cortex in pSTG/S

In our primary analysis, we focused on regions of the pSTG/S that preferred visual presentation of faces making mouth movements to faces making eye movements. To determine if the observed effects were specific to mouth-preferring pSTG/S, in a control analysis we considered eye-preferring pSTG/S. Voxels within each hemisphere meeting three criteria (anatomical location in the pSTG/S/STG; significant BOLD response to visual stimulation compared with fixation baseline; and a greater response to eye movements) were grouped into an ROI. pSTG/S eye ROIs were successfully created in each of the 44 hemispheres that were examined (see Figure S1 for ROIs in 8 sample hemispheres). For visualization only, a group map of pSTG/S eye-preferring regions was created on the cortical surface (Figure 6A).

**Figure 6:**
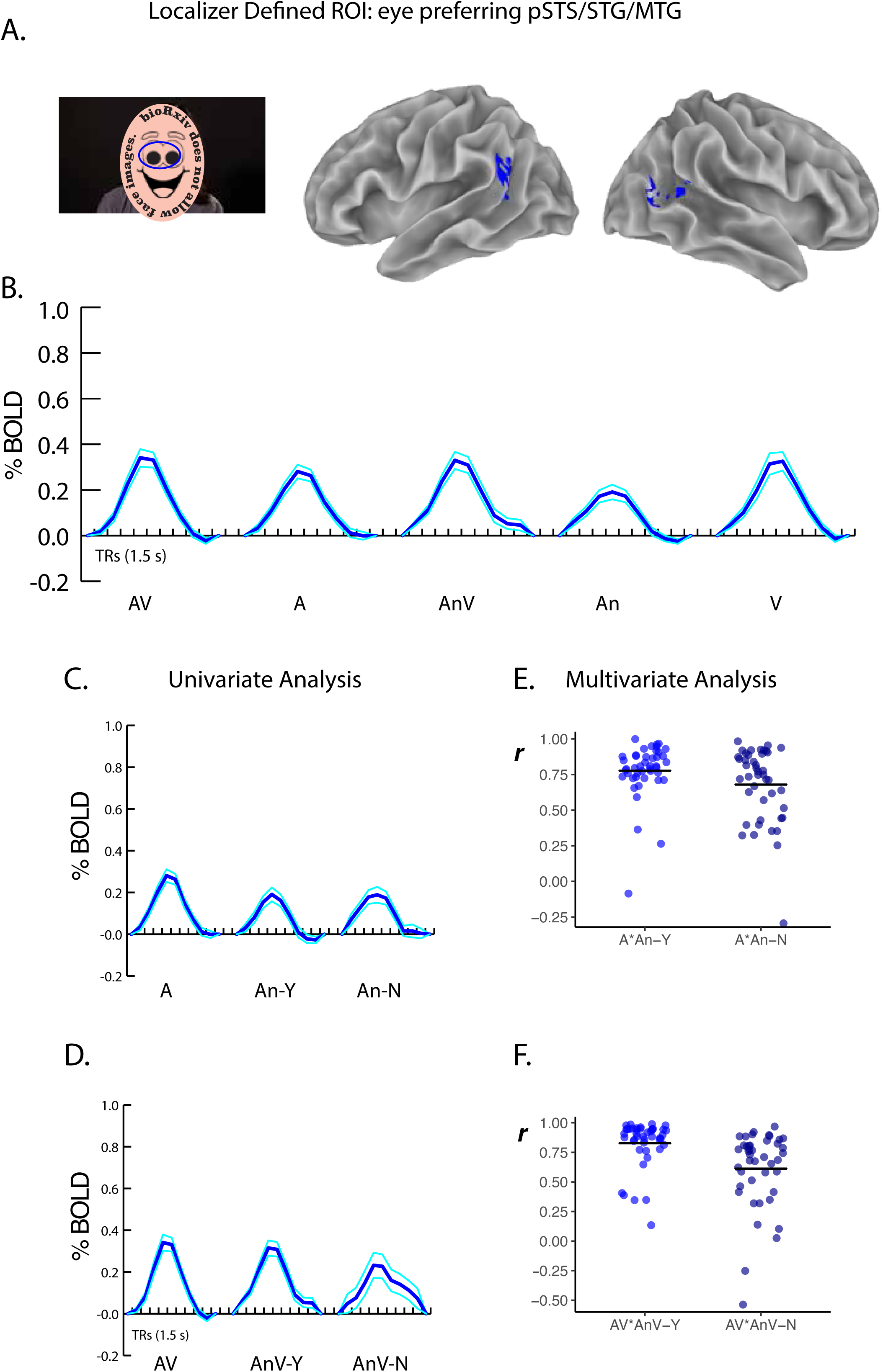
**A**. In a localizer fMRI experiment, silent videos of facial eye and mouth movements were presented. Image shows still frame from silent video of eye movements, blue ellipse highlights eye movement. In each individual participant, eye-preferring regions in the left and right pSTG/S were identified. For visualization only, a group map was created (blue color; Fig. S1 shows ROIs in individual participants). **B**. For each participant, the time course of the BOLD fMRI responses to the five different sentence types was calculated within the eye-preferring ROI. Thick blue lines show mean across participants, thin cyan lines show standard error of the mean. **C**. For each participant, the time course of the BOLD fMRI response to noisy auditory speech rated as intelligible (An-Y) and noisy auditory speech rated as not intelligible (An-N) in the eye-preferring ROI was calculated (response to A speech shown for reference). Thick red lines show mean across participants, thin orange lines show standard error of the mean. **D**. For each participant, the time course of the response to noisy audiovisual speech rated as intelligible (AnV-Y) and noisy auditory speech rated as not intelligible (AnV-N) in the eye-preferring ROI was calculated (response to AV speech shown for reference). **E**. Within subject activation pattern correlations in the eye-preferring ROI between clear auditory (A) and noisy auditory speech rated as intelligible (An-Y) (left points) and clear auditory (A) and noisy auditory speech rated as not intelligible (An-N) (right points). **F**. Within subject activation pattern correlations between clear audiovisual (AV) and noisy audiovisual speech rated as intelligible (AnV-Y) (left points) and clear audiovisual (AV) and noisy audiovisual speech rated as not intelligible (AnV-N) (right points).

Mouth-preferring voxels were located more anteriorly in the pSTG/S, while eye-preferring voxels were located more posteriorly. To quantify this effect, we calculated the center of mass of the activation for the mouth and eye ROIs. The average Euclidean distance between the centers of mass of the mouth and eye ROIs was 15 ± 5 mm (SD) in standard space in the left hemisphere (*t*-test against zero: *t*_(21)_ = 12, *p* = 9.5 × 10^−11^) and 14 ± 7 mm in the right hemisphere (*t*-test against zero: *t*_(21)_ = 10.5, *p* = 7.8 × 10^−10^). The primary driver of this effect was a more anterior location for mouth-preferring voxels (results for all cardinal axes shown in Table 1). There was no significant difference in volume of activation between the mouth and eye ROIs in either the left hemisphere (average volume: 2600 ± 2100 mm^3^ for eye and 2200 ± 2100 mm^3^ for mouth, *t*-test, *t*_(21)_ = 0.48, *p* = 0.64) or the right hemisphere (3200 ± 2300 mm^3^ for mouth and 1900 ± 1900 mm^3^ for eye, *t*_(21)_ = 1.6, *p* = 0.13).

**Table 1.** ROI sizes and location.

| pSTG/S | Size (mm<sup>3</sup>) | Talairach coordinates (mm) and statistical difference |  |  |  |  |  |
| --- | --- | --- | --- | --- | --- | --- | --- |
|  |  | x | *t, p* | y | *t, p* | z | *t, p* |
| Left |  |  |  |  |  |  |  |
| Mouth | 2575 ± 2074 | **−56 ± 3** | 4.20 | **−44 ± 6** | 4.86 | 10 ± 3 | 1.58 |
| Eye | 2194 ± 2111 | **−51 ± 4** | 4.2×10<sup>−4</sup> | **−53 ± 7** | 9.4×10<sup>−5</sup> | 12 ± 5 | 0.131 |
| Right |  |  |  |  |  |  |  |
| Mouth | 3178 ± 2303 | **54 ± 3** | 3.47 | **−42 ± 5** | 5.95 | **8 ± 3** | 3.871 |
| Eye | 1916 ± 1920 | **51 ± 4** | 2.3×10<sup>−3</sup> | **−52 ± 5** | 7.3×10<sup>−6</sup> | **14 ± 6** | 8.8×10<sup>−4</sup> |

### Univariate responses in pSTG/S eye ROIs

Unlike in pSTG/S mouth ROIs, in pSTG/S eye ROIs there were no significant differences between AV and A speech (0.30% vs. 0.25%; *χ*^*2*^_(1)_ = 2.04, *p* = 0.153); between AV and V speech (0.30% vs. 0.29%; *χ*^*2*^_(1)_ = 0.04, *p* = 0.842); between A and V speech (0.25% vs. 0.29%; *χ*^*2*^_(1)_ = 0.96, *p* = 0.328; see Figure 6B); between intelligible and unintelligible noisy auditory sentences (An-Y *vs*. An-N, 0.17% vs. 0.17%; χ^*2*^_(1)_ = 0.0, *p* = 0.95; Figure 6C); or between intelligible and unintelligible noisy audiovisual sentences (AnV-Y *vs*. AnV-N, 0.29% *vs*. 0.30%, *χ*^*2*^_(1)_ = 0.06, *p* = 0.81; Figure 6D).

### Multivariate responses in pSTG/S eye ROIs

For auditory-only sentences, the response patterns were more similar between clear and intelligible noisy sentences than between clear and unintelligible noisy sentences (*r*_A*An-Y_ *=* 0.78 *vs. r*_A*An-N_ *=* 0.68, *χ*^*2*^_(1)_ = 5.37, *p* = 0.02, Figure 6E). The same was true for audiovisual sentences (*r*_AV*AnV-Y_ = 0.83 *vs. r*_AV*AnV-N_ = 0.59, 0.83 *vs*. 0.59, *χ*^*2*^_(1)_ = 29, *p* = 8.8 × 10^−8^, Figure 6F). The linear mixed-effects model found no significant main effect for stimulus type (*χ*^*2*^_(1)_ = 0.41, *p* = 0.52) but a significant effect for percept (*χ*^*2*^_(1)_ = 28, *p* = 6.4 × 10^−7^) and a significant interaction (*χ*^*2*^_(1)_ = 4.6, *p* = 0.03).

### Univariate comparison of pSTG/S mouth and eye ROIs

Mouth-preferring pSTG/S showed significantly greater responses to AV than A or V sentences while eye-preferring pSTG/S did not. To determine if this difference was itself significant, a linear mixed-effects models was fit with fixed factors of ROI and physical stimulus condition (A, An, AV, AnV, V). There were significant main effects for ROI (*χ*^*2*^_(1)_ = 36, *p* = 2.3 × 10^−9^) and speech type (*χ*^*2*^_(6)_ = 90, *p* = < 2.2 × 10^−16^) as well as a significant interaction between ROI and condition (*χ*^*2*^_(6)_ = 37, *p* = 1.8 × 10^−7^). Complete results in Table 2.

**Table 2:** Analyses of responses to different speech types without behavior in mouth and eye ROIs.

|  | Estimate | Std. Error | *df* | *t* | *p* |
| --- | --- | --- | --- | --- | --- |
| **Fixed effects** |  |  |  |  |  |
| A (Intercept) | 0.25 | 0.04 | 52.7 | 6.32 | $5.8 \times 10^{-8}$ |
| AN | -0.09 | 0.04 | 396 | -2.26 | 0.024 |
| AV | 0.05 | 0.04 | 396 | 1.32 | 0.187 |
| ANV | 0.05 | 0.04 | 396 | 1.33 | 0.184 |
| V | 0.04 | 0.04 | 396 | 1.10 | 0.273 |
| Mouth | 0.35 | 0.07 | 39.6 | 5.26 | $5.3 \times 10^{-6}$ |
| AN × Mouth | -0.04 | 0.05 | 396 | -0.71 | 0.478 |
| AV × Mouth | 0.10 | 0.05 | 396 | 1.90 | 0.058 |
| ANV × Mouth | 0.09 | 0.05 | 396 | 1.61 | 0.107 |
| **Random effects** |  |  |  |  |  |
|  | Variance |  |  |  |  |
| Participant | 0.019 |  |  |  |  |
| Participant × ROI | 0.065 | | | | |

**Table 2 Legend:** We used a functional localizer to define mouth and eye selective regions of the pSTG/S.
| *Main effects* | *df* | $\chi^2$ | *p* |
| --- | --- | --- | --- |
| ROI | 1 | 35.71 | $2.3 \times 10^{-9}$ |
| Speech Type | 4 | 89.54 | $< 2.2 \times 10^{-16}$ |
| *Interaction* |  |  |  |
| ROI × Speech type | 4 | 37.05 | $1.8 \times 10^{-7}$ |

Mouth-preferring pSTG/S showed significantly greater responses to intelligible *vs*. unintelligible sentences; eye-preferring pSTG/S did not. A linear mixed-effects model was fit with factors ROI (mouth pSTG/S, eye pSTG/S) and stimulus condition split by perception (A, An-Y, An-N, AV, AnV-Y, AnV-N, V). There were significant main effects for ROI (*χ*^*2*^_(1)_ = 37, *p* = 9.3 × 10^−10^) and speech type (*χ*^*2*^_(6)_ = 98, *p* = < 2.2 × 10^−16^) as well as a significant interaction between ROI and condition (*χ*^*2*^_(6)_ = 34, *p* = 6.4 × 10^−6^). Complete results in Table 3.

**Table 3:**
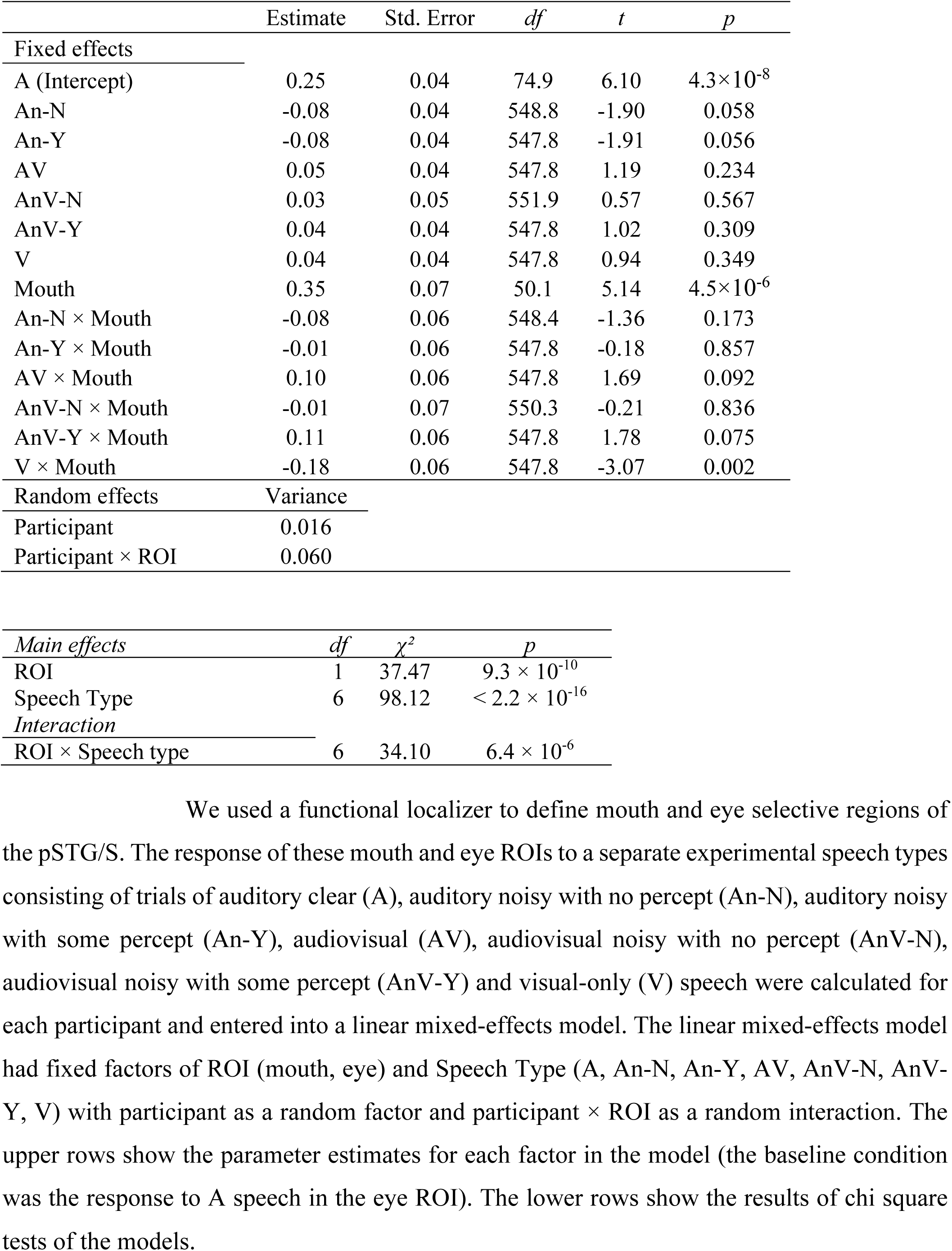
Analyses of responses to different speech types including behavior in mouth and eye ROIs.

|  | Estimate | Std. Error | *df* | *t* | *p* |
| --- | --- | --- | --- | --- | --- |
| **Fixed effects** |  |  |  |  |  |
| A (Intercept) | 0.25 | 0.04 | 74.9 | 6.10 | $4.3 \times 10^{-8}$ |
| An-N | -0.08 | 0.04 | 548.8 | -1.90 | 0.058 |
| An-Y | -0.08 | 0.04 | 547.8 | -1.91 | 0.056 |
| AV | 0.05 | 0.04 | 547.8 | 1.19 | 0.234 |
| AnV-N | 0.03 | 0.05 | 551.9 | 0.57 | 0.567 |
| AnV-Y | 0.04 | 0.04 | 547.8 | 1.02 | 0.309 |
| V | 0.04 | 0.04 | 547.8 | 0.94 | 0.349 |
| Mouth | 0.35 | 0.07 | 50.1 | 5.14 | $4.5 \times 10^{-6}$ |
| An-N × Mouth | -0.08 | 0.06 | 548.4 | -1.36 | 0.173 |
| An-Y × Mouth | -0.01 | 0.06 | 547.8 | -0.18 | 0.857 |
| AV × Mouth | 0.10 | 0.06 | 547.8 | 1.69 | 0.092 |
| AnV-N × Mouth | -0.01 | 0.07 | 550.3 | -0.21 | 0.836 |
| AnV-Y × Mouth | 0.11 | 0.06 | 547.8 | 1.78 | 0.075 |
| V × Mouth | -0.18 | 0.06 | 547.8 | -3.07 | 0.002 |
| **Random effects** |  |  |  |  |  |
|  | Variance |  |  |  |  |
| Participant | 0.016 |  |  |  |  |
| Participant × ROI | 0.060 | | | | |

**Table 3 Legend:** We used a functional localizer to define mouth and eye selective regions of the pSTG/S.
|  |  |  |  |
| --- | --- | --- | --- |
| *Main effects* | *df* | $\chi^2$ | *p* |
| ROI | 1 | 37.47 | $9.3 \times 10^{-10}$ |
| Speech Type | 6 | 98.12 | $< 2.2 \times 10^{-16}$ |
| *Interaction* |  |  |  |
| ROI × Speech type | 6 | 34.10 | $6.4 \times 10^{-6}$ |

### Multivariate comparison of pSTG/S mouth and eye ROIs

Response patterns for the different kinds of sentences were more similar in mouth-preferring pSTG/S than in eye-preferring pSTG/S (Figure 7A) with mean *r* of 0.70 *vs*. 0.66 averaged over all pair-wise stimulus correlations, paired *t* test *t*_(20)_ = 4.1, *p* = 6.1 × 10^−4^. Comparing conditions, intelligible sentences (A, An-Y, AV, AnV-Y) evoked the most similar response patterns, and this correlation was higher in mouth-preferring than eye-preferring pSTG/S, mean *r* of 0.86 *vs*. 0.76 averaged over pair-wise correlations within this subset of sentences, paired *t* test: *t*_(42)_ = 3.0, *p* = 0.005.

**Figure 7:**
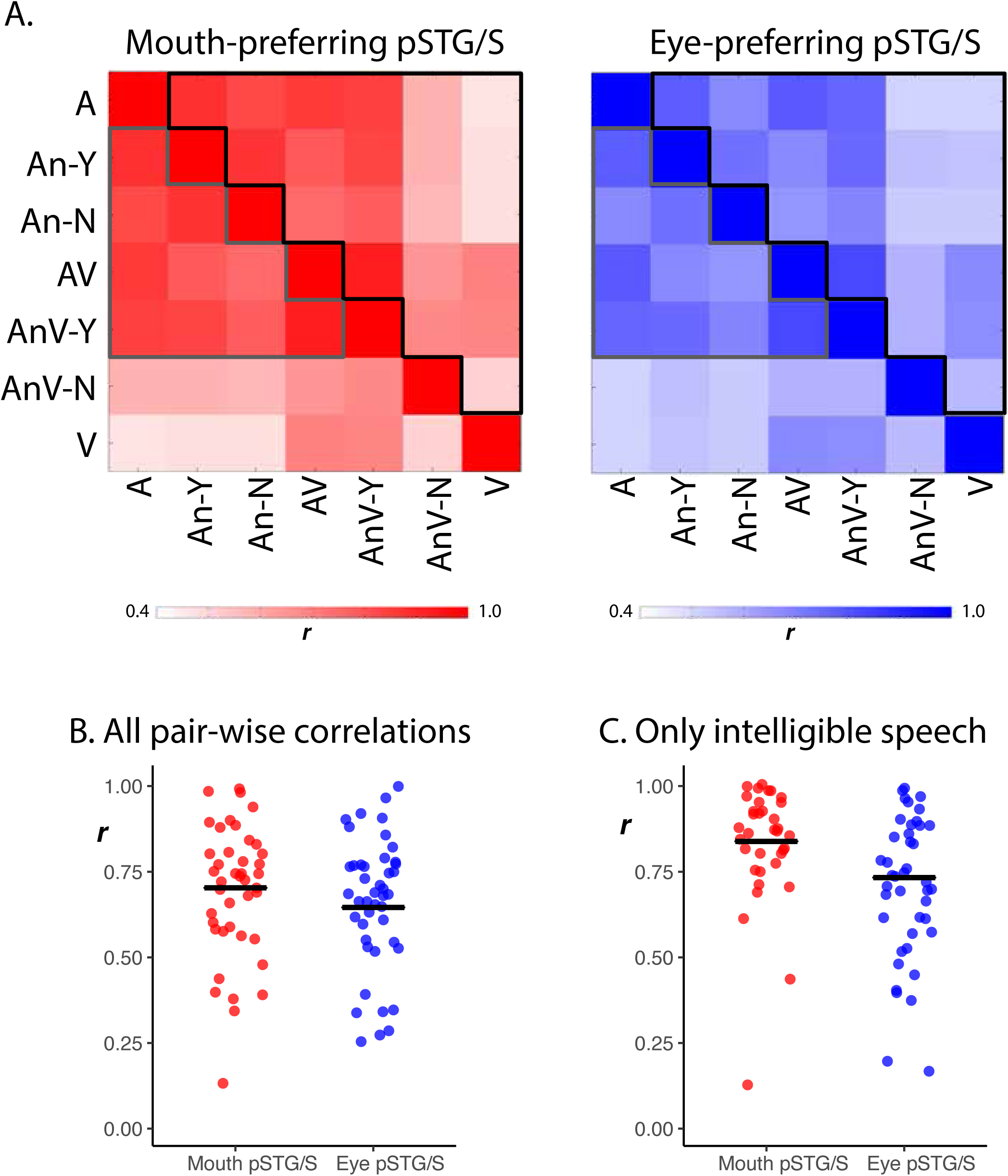
**A**. Average correlation matrix between all speech conditions in the mouth-preferring pSTG/S (left panel, reproduced from Figure 5A) and eye-preferring pSTG/S (right panel). For each participant, the activation patterns were correlated between every pair of conditions to create a matrix of correlation values. These matrices were then averaged across all participants. The black box highlights pairwise correlations calculated for individual participants, shown in panel B. The gray box highlights pairwise correlations calculated for individual participants shown in panel C. **B**. Individual participant correlation values, one dot per participant. Within each participant, all correlation values from the correlation matrix were averaged, black lines show group mean. **C**. Individual participant correlation values, one dot per participant. Within each participant, only pair-wise correlation values from sentences in which the speech was reported as intelligible were averaged (AV, A, An-Y, AnV-Y).

## Discussion

We used a localizer to identify subregions of the pSTG/S selective for the visual mouth movements that comprise visual speech in individual participants (13, 17). A control ROI consisted of pSTG/S subregions selective for viewed eye movements. In the main experiment, we measured responses in both subregions of the pSTG/S to clear and noisy auditory-only and audiovisual sentences. As expected, adding visual speech to noisy auditory speech greatly improved intelligibility (reviewed in 1). *Post hoc* trial sorting was used to measure BOLD fMRI responses to sentences that were more or less intelligible.

In the univariate analysis, we observed stronger responses for clear auditory-only speech *vs*. noisy auditory-only speech in both mouth-preferring and eye-preferring subregions of pSTG/S, consistent with many previous reports of stronger BOLD signals for clear speech throughout lateral temporal cortex (9, 21-23).

We observed significantly stronger responses for intelligible *vs*. unintelligible speech (both auditory-only and audiovisual) in mouth-preferring but not eye-preferring pSTG/S, and this difference between subregions was significant when tested with a linear mixed-effects model. Most previous studies used volumetric group analysis, blurring together mouth and eye-preferring regions due to anatomical variability between participants. This may partially explain why Bishop and Miller (9) did not to find an effect of intelligibility in pSTG/S. In support of this idea of anatomical heterogeneity, Evans and colleagues described a small region in mid-posterior STG that was positively correlated with a post-scanning behavioral task across participants (23).

In addition to the univariate analysis, the localizer allowed for an examination of the multivariate pattern of activity across the population of voxels in the mouth-preferring pSTG/S subregion. The largest differences between response patterns were between sentences with different sensory components, such as the responses to auditory-only and visual-only speech. However, there was also a large difference in response patterns between intelligible and unintelligible speech (with sensory information held constant). Auditory-only noisy speech that was intelligible evoked a response pattern more similar to clear auditory speech (*r*_A*An-Y_ = 0.88 *vs. r*_A*An-N_ = 0.83). This effect was even larger for noisy audiovisual sentences. For noisy audiovisual sentences that were understood, the pattern of activity was very similar to that for clear audiovisual sentences with no added noise (*r*_AV*AnV-Y_ = 0.93). In contrast, for unintelligible sentences, the pattern of activity was very different (*r*_AV*AnV-N_ = 0.65). Even though the physical stimuli were similar (videos of talking faces + noisy auditory speech), the pronounced difference in response patterns suggests that successful integration of visual and noisy auditory speech normalized the response pattern, resulting in accurate perception. Taken together, these results suggest that the sensory input into pSTG/S (auditory-only *vs*. visual-only speech), the intelligibility of speech, and successful multisensory integration of auditory and visual speech components are all important contributors to the pSTG/S response. Consistent with our results, Tuennerhoff and Noppeney reported that responses in pSTG/S are driven both by the physical stimulus and the resulting percept (24).

An important question for further research is determining the mechanisms by which visual speech modulates responses to auditory speech in pSTG/S. One possibility is that the early arrival of visual information influences the responses of populations of neurons in pSTG/S selective for different speech sounds, exciting populations that are compatible with the viewed mouth movements and inhibiting populations that are incompatible with it (6). From a practical perspective, normalizing the response pattern in the service of better comprehension could be accomplished through real-time fMRI neurofeedback (25) or through brain-computer interfaces (26).

### Comparison with previous localizer results

The freely-available stimuli available in the pSTG/S localizer of (13) were used to identify mouth-preferring and eye-preferring regions of pSTG/S. The present study replicated the main findings of (13) in a different group of participants. First, mouth-preferring and eye-preferring regions of pSTG/S were identified in every participant. Second, mouth-preferring regions responded significantly more to auditory-only speech than eye-preferring regions even though the localizer contained only visual stimuli. The present result extends the original study by showing that mouth-preferring subregions, but not eye-preferring subregions, demonstrate the characteristic neuroimaging signature of multisensory integration, with a significantly greater response for audiovisual speech than auditory-only or visual-only speech (27). The present study also adds support for the idea that the mouth-preferring subdivision has a special role in multisensory speech perception, since it responded differently (in both univariate and multivariate analyses) when speech was intelligible than when it was not. Eye and mouth-preferring regions of the pSTG/S can be identified during free viewing of natural speech in which fixations of the mouth region last only a few hundred milliseconds (17). Therefore, techniques with faster temporal resolution than BOLD fMRI, such as MEG (28), EEG (29) and iEEG (30) will be important in further elucidating the multisensory computations performed by pSTG/S.

## Acknowledgements

This work was supported by the National Institutes of Health (R01NS065395 to M.S.B) and the Deutsche Forschungsgemeinschaft (RE 3693/1-1 to J.R.). We acknowledge the Core for Advanced MRI at Baylor College of Medicine and Kira Wegner-Clemens for assistance with MR scanning.

The authors declare no competing financial interests.

## Figure captions

**Figure S1:**
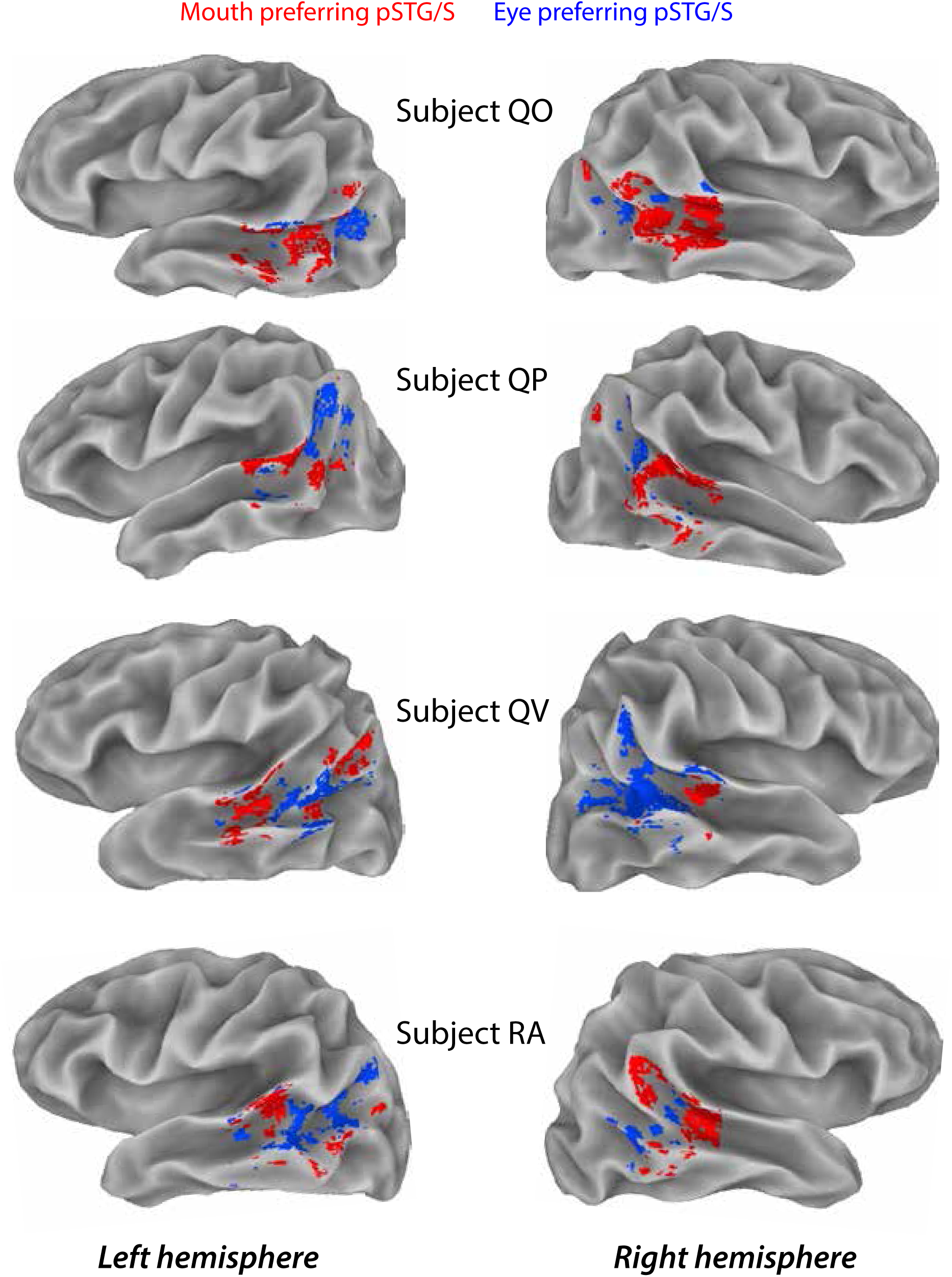
Single participant activation maps showing mouth-preferring (red) and eye-preferring (blue) subregions of pSTG/S.

